# Identification of transcriptional landscapes and functions of the inhibition of ARHGEF2 in hepatocellular carcinoma cells

**DOI:** 10.1101/2022.08.18.504433

**Authors:** Min Zhang, David C He, Andrew Chung

## Abstract

The RHO guanine exchange factor ARHGEF2 has exchange activity toward RHOA, which is essential for the development of cancers such as liver cancer. However, the potential functions and mechanisms of ARHGEF2 in the progression of liver cancer are largely unknown. In this study, we identified the transcriptional landscapes of hepatocellular carcinoma cells treated with ARHGEF2 shRNAs. The gene enrichment assays such as KEGG and GO were used to further analyze the potential signaling pathways. Moreover, the PPI network and Reactome map were used to further identify the biological processes. The results showed that Alzheimer’s disease disease (AD) and Cushing syndrome (CS) are the major signaling pathways involved in the ARHGEF2-shRNAs treated hepatocellular carcinoma cells. We identified the top ten interactive genes including ICAM1, APOE, LDLR, NAT10, HSPA1A, EDN1, CACNA1C, KCNMA1, SNAI1, and ELN. Our study may provide novel mechanisms for the treatment of liver cancer by inhibiting ARHGEF2.

## Introduction

Hepatocellular carcinoma is a primary malignancy of the liver and is the most common cause of death from cancer in some countries in the world^1^. Although it is less common in most parts of the developed western world, it shows to be increasing substantially in incidence^2^. It usually happens in the setting of chronic liver disease, and the diagnosis of hepatocellular carcinoma is often performed by gastroenterologists and hepatologists^3^. It was reported an increase of approximately 80% in the incidence of HCC in the USA over the past decades and it was estimated that about 15,000 new cases per year^4^. The reasons for the increase are not sure but it has contributed to the occurrence of hepatitis C and B-related HCC^5^.

Rho guanine nucleotide factor 2 is an important member of the RhoGEF family^6^. Rho family members contain RhoA, RhoB, and RhoC, and multiple RAC and CDC42 isoforms^7^. Rho GTPases control the cellular cytoskeleton and cell survival by mediating the activity of Rho family members^8^. RhoA activation is necessary for transformation by oncogenic RAS via sustained mitogen-activated protein kinase (MAPK) signaling to increase proliferation and migration^9^. ARHGEF2 is critical for the survival and growth of RAS-transformed cells^10^. ARHGEF2 is a microtubule-associated Dbl-related GEF with typical exchange activity for RhoA^11^. Studies showed that amplified ARHGEF2 promotes cell motility by activating RhoA signaling, suggesting that ARHGEF acts as a therapeutic target in several types of cancers, including hepatocellular cancer^12^. However, the mechanism of the transcriptional regulation of ARHGEF2 in liver cancer is not clear.

Our aim is to identify the significant signaling pathways and molecules by analyzing the RNA-seq data from the hepatocellular carcinoma treated by ARHGEF2 shRNAs. We also performed the KEGG, GO, Reactome map, and PPI to further study the potential functions of each key molecule. Therefore, our study may provide the functional mechanisms in the treatment of hepatocellular carcinoma by inhibiting ARHGEF2.

## Materials and Methods

### Acquisition of microarray data

GEO database (http://www.ncbi.nlm.nih.gov/geo/) was used for downloading the following gene expression profile dataset: GSE205757. Illumina HiSeq 4000 (Homo sapiens) (The First Affiliated Hospital of Anhui Medical University, The First Affiliated Hospital of Anhui Medical University, Hefei, Anhui 230000, China). The dataset was chosen based on the criteria as follows: Studies comparing control shRNAs treated HepG2 cells with ARHGEF2 shRNAs treated HepG2 cells; Studies included gene expression profiles; Datasets were excluded if the study did not contain control groups; Datasets from other organisms were excluded.

### Identification of differentially expressed genes (DEGs)

The gene expression matrix data in the GSE205757 dataset was normalized and processed through the R program as described^13–18^. Then, the empirical Bayesian algorithm in the “limma” package was applied and compared between patients with diseases and the healthy control population as controls with the detection thresholds of adjusted P-values <0.05 and |log2FC| >1.

### Gene intersection between DEGs

We intersected the DEGs screened by the “limma” package with the modular genes. The intersecting genes were analyzed for Kyoto Encyclopedia of Genes and Genomes (KEGG) and Gene Ontology (GO) enrichment using the “clusterprofiler” package in the R program. The results were then visualized using the “ggplot2” package.

### Protein-protein interaction (PPI) networks

The interacting genes were constructed using Protein-Protein Interaction Networks Functional Enrichment Analysis (STRING; http://string.db.org). The Molecular Complex Detection (MCODE) was used to analyze the PPI networks. This method predicts the PPI network and provides an insight into the biological mechanisms. The biological processes analyses were further performed by using Reactome (https://reactome.org/), and P<0.05 was considered significant.

## Results

### DEGs in hepatocellular carcinoma HepG2 cells with the ARHGEF2 knockdown

To make sure the effects of ARHGEF2 knockdown on hepatocellular carcinoma cells, we used the RNA-seq data from the GEO database (GSE205757). We identified the 511 significantly changed genes (P < 0.01). The significant increased and decreased genes were also indicated by the heatmap (Figure 1). We further presented the top ten significant genes in Table 1.

**Figure 1.**
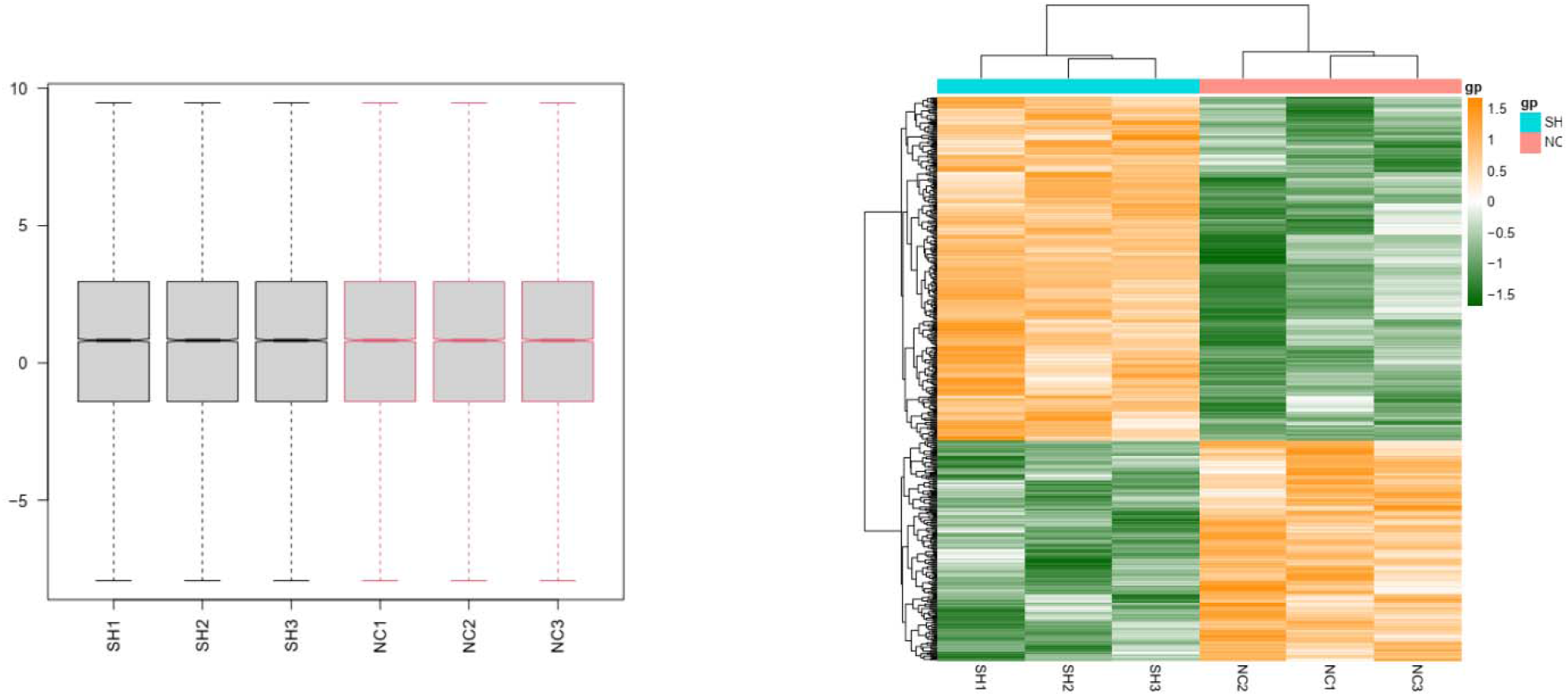
Heatmap in hepatocellular carcinoma HepG2 cells with the ARHGEF2 knockdown. (A) DEGs (P < 0.05) were used to construct the heatmap. NC, normal control shRNA; SH, ARHGEF2 shRNA.

**Table 1.**
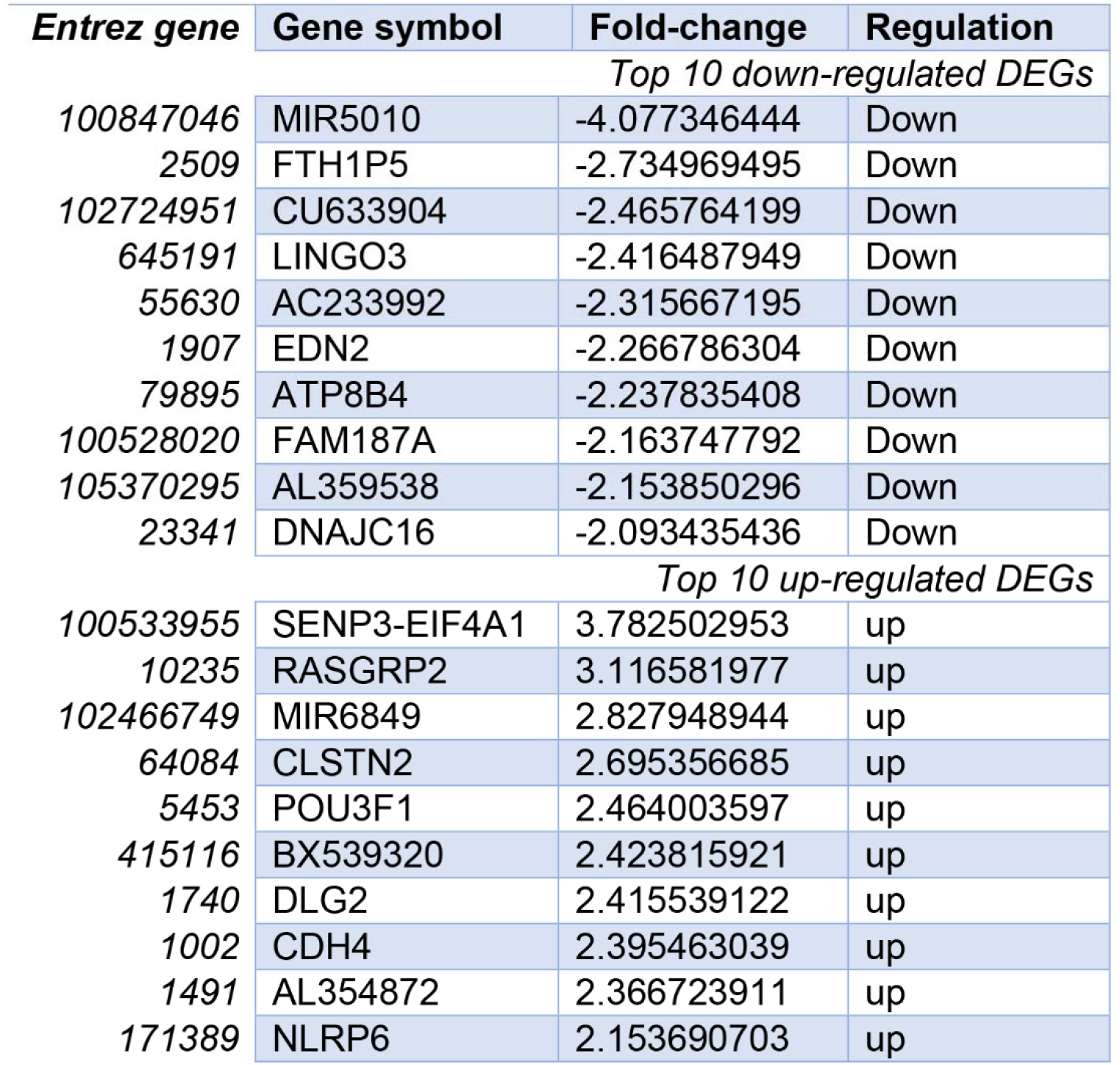

### Gene enrichment and function analyses in hepatocellular carcinoma HepG2 cells with the ARHGEF2 knockdown

To study the mechanisms in hepatocellular carcinoma HepG2 cells with the ARHGEF2 knockdown, we performed the gene enrichment analysis (Figure 2). We found the top changed KEGG signaling pathways: “Alzheimer disease”, “Cushing syndrome”, “Hippo signaling pathway”, “Axon guidance”, “Toxoplasmosis”, “Vascular smooth muscle contraction”, “Cardiac muscle contraction”, “ECM-receptor interaction”, “Hypertrophic cardiomyopathy”, “Cholesterol metabolism”. We found the top changed Biological Processes: “axon guidance”, “neuron projection guidance”, “regulation of tube diameter”, “blood vessel diameter maintenance”, “regulation of tube size”, “microtubule polymerization or depolymerization”, “Rho protein signal transduction”, “platelet-derived growth factor receptor signaling pathway”, “positive regulation of steroid metabolic process”, “regulation of ketone biosynthetic process”. We figured out the significant changed cellular components: “collagen-containing extracellular matrix”, “endoplasmic reticulum lumen”, “basement membrane”, “clathrin-coated vesicle membrane”, “neuromuscular junction”, “specific granule membrane”, “clathrin–coated endocytic vesicle”, “aggresome”, “laminin complex”, “synaptic cleft”. We also figured out the top changed molecular functions: “GTPase activity”, “microtubule binding”, “ATP hydrolysis activity”, “extracellular matrix binding”, “laminin binding”, “insulin-like growth factor binding”, “lipoprotein particle binding”, “protein-lipid complex binding”, “frizzled binding”, “neurotransmitter binding”.

**Figure 2.**
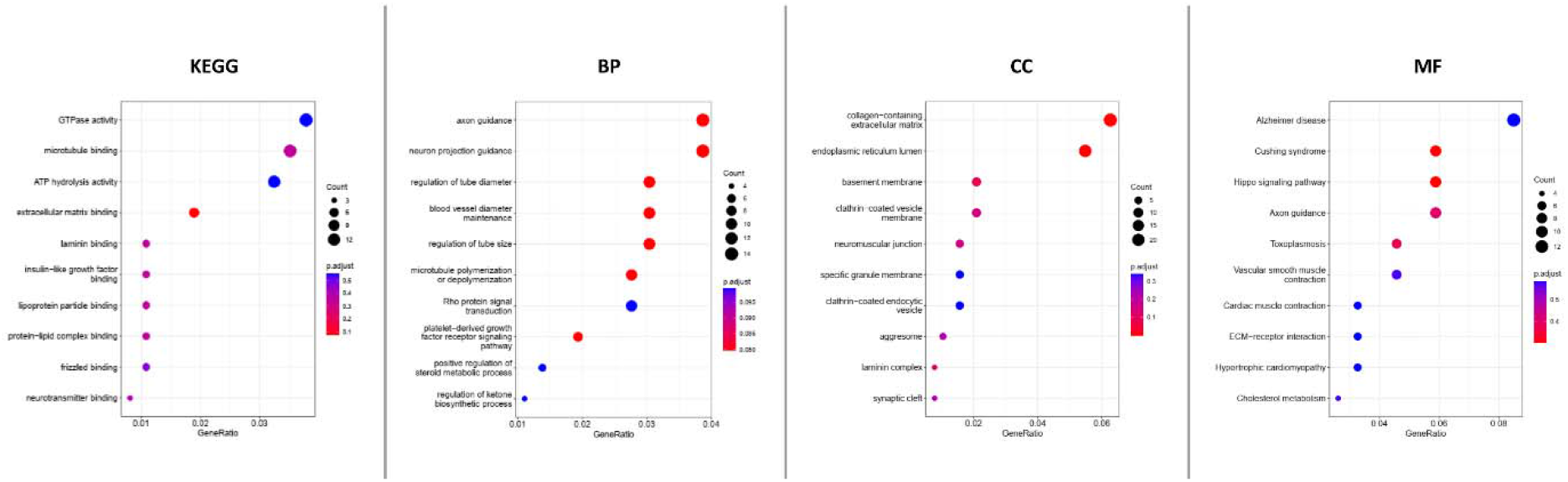
KEGG and GO analyses of DEGs in hepatocellular carcinoma HepG2 cells with the ARHGEF2 knockdown. KEGG analysis; BP: Biological processes; CC: Cellular components; MF: Molecular functions.

### The protein-protein interaction (PPI) network and Reactome analyses

The PPI network was created by using 432 nodes and 454 edges with the Cytoscape software. The top ten genes with the highest degree scores were shown in Table 2. By analyzing the PPI network, we further constructed the top two clusters in Figure 3. We then constructed the Reactome map (Figure 4) and performed the Reactome analysis, which includes “Attenuation phase”, “HSF1-dependent transactivation”, “HSF1 activation”, “Regulation of HSF1-mediated heat shock response”, “Cellular response to heat stress”, “Netrin mediated repulsion signals”, “TFAP2 (AP-2) family regulates transcription of growth factors and their receptors”, “Transcriptional regulation by the AP-2 (TFAP2) family of transcription factors”, “FLT3 signaling through SRC family kinases”, and “TNFR1-mediated ceramide production” (Supplemental Table S1).

**Table 2.** Top ten genes demonstrated by connectivity degree.

| **Gene symbol** | **Gene title** | **Degree** |
| --- | --- | --- |
| *ICAM1* | Intercellular Adhesion Molecule 1 | 16 |
| *APOE* | Apolipoprotein E | 15 |
| *LDLR* | Low Density Lipoprotein Receptor | 14 |
| *NAT10* | N-Acetyltransferase 10 | 14 |
| *HSPA1A* | Heat Shock Protein Family A (Hsp70)<br>Member 1A | 11 |
| *EDN1* | Endothelin 1 | 11 |
| *CACNA1C* | Calcium Voltage-Gated Channel Subunit<br>Alpha1 C | 11 |
| *KCNMA1* | Potassium Calcium-Activated Channel<br>Subfamily M Alpha 1 | 10 |
| *SNAI1* | Snail Family Transcriptional Repressor 1 | 9 |
| *ELN* | Elastin | 9 |

**Figure 3.**
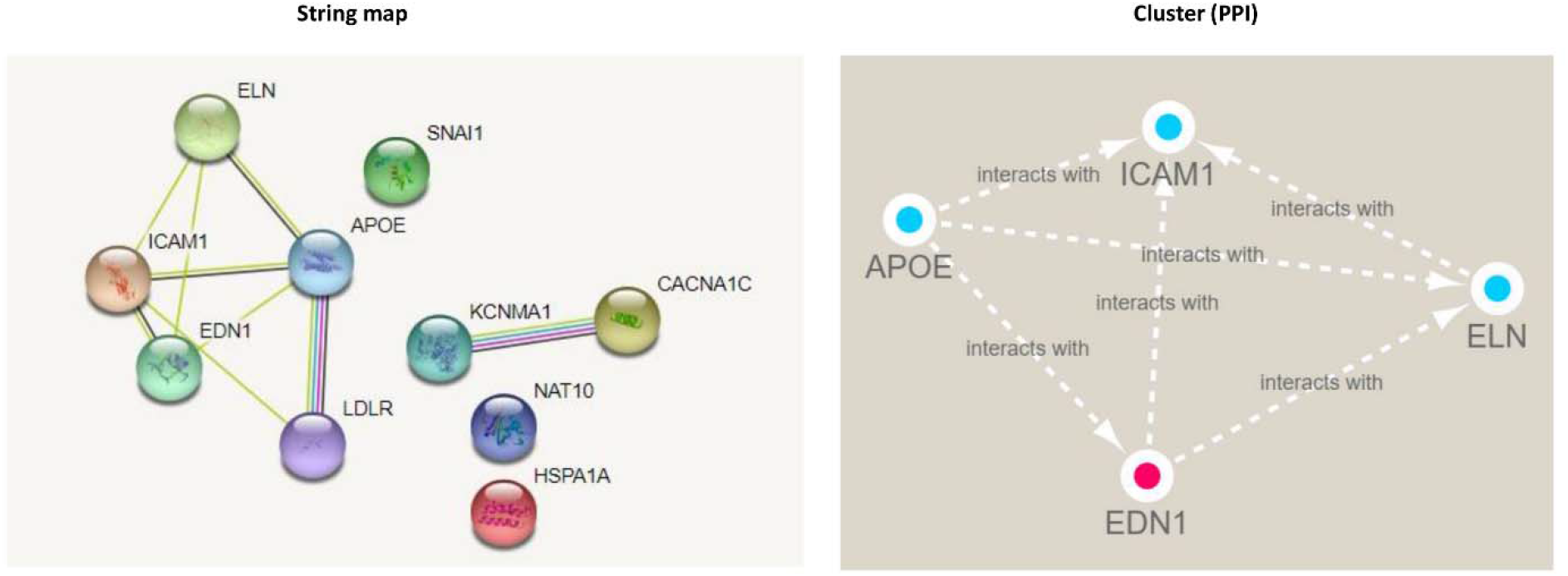
The PPI network analyses of DEGs in hepatocellular carcinoma HepG2 cells with the ARHGEF2 knockdown. The Cluster map were constructed according to the String map by MCODE.

**Figure 4.**
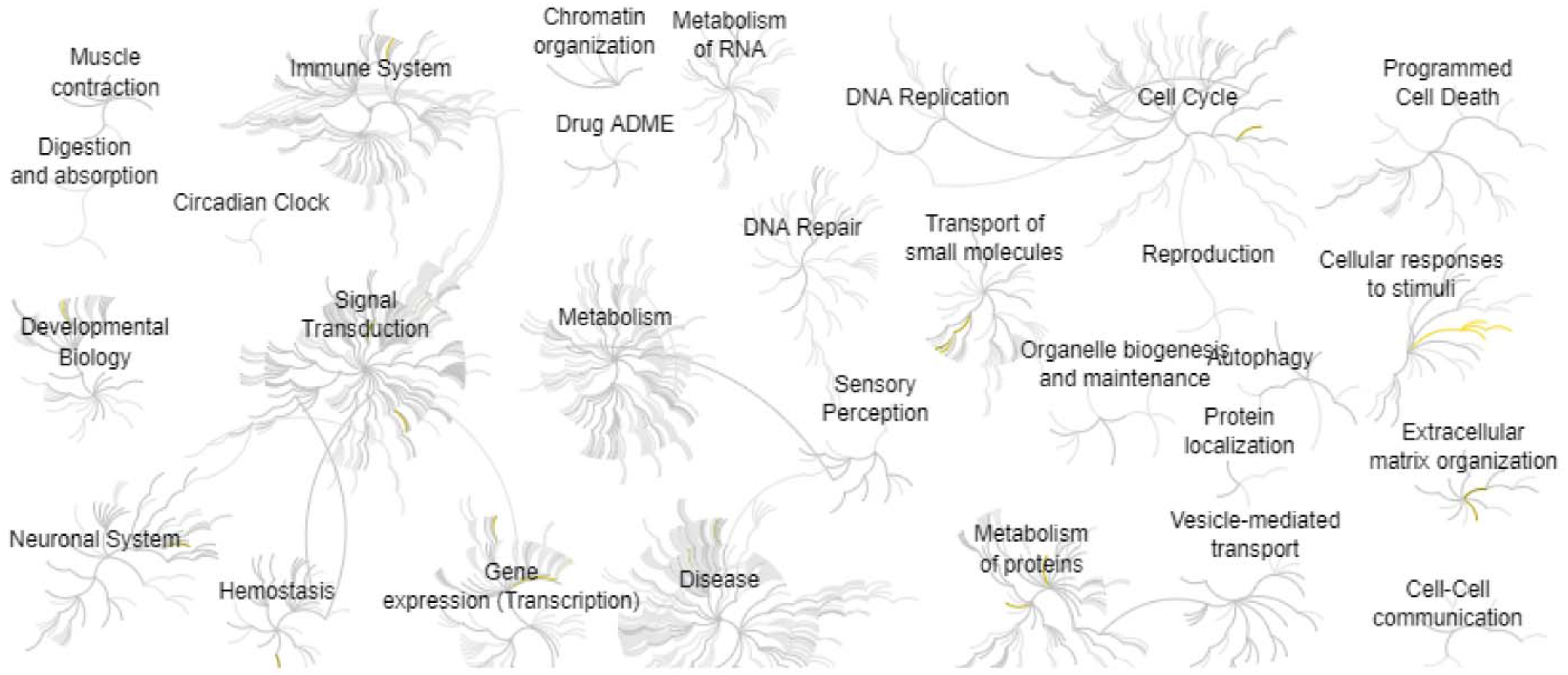
Reactome map representation of the significant biological processes in hepatocellular carcinoma HepG2 cells with the ARHGEF2 knockdown.

## Discussion

Liver cancer is a common malignancy for prevention and treatment. Moreover, the mechanism of the liver cell progression is still unknown^19^. ARHGEF2 is a RhoA activator and acts as a key downstream target of YTHDF1, which is critical for the progression of colorectal cancer^20^. Our study further identifies the key molecules after inhibition of ARHGEF2 in hepatocellular carcinoma by using the bioinformatic method.

By performing the gene enrichment, we identified the top two related signaling pathways, including “Alzheimer’s disease (AD)” and “Cushing syndrome (CS)”. Interestingly, Kelly N H Nudelman et al also found several biological alterations of cancer cells between AD and cancer, including increased inflammation, increased cell death, and decreased suppressors^21^. Constantine A Stratakis et al found that adrenocortical tumors are related to CS which showed the inhibited cAMP signaling pathway, but adrenal cancer is associated with decreased expression of growth factors and germline mutations of tumor suppressor genes including TP53^22^.

By constructing the PPI network, we identified the top key molecules affected by ARHGEF2 in hepatocellular carcinoma (Table 2). The study by Stine L Figenschau et al showed that the expression of ICAM1 is induced by inflammatory cytokines and related to TLS formation in breast cancers^23^. Circadian gene clocks regulate gene expression through translational and post-translational regulations to maintain cell functions^24–34^. CAM1 acts as a key circadian regulated factor that can promote the adhesion of monocytes and endothelial cells^35^. The study by Masoud F Tavazoie et al found that ApoE controls innate immune suppression and acts as a target for increasing the efficacy of cancer immunotherapy in patients^36^. Ziye Chen et al found the inhibition of LDLR can increase proliferation and metastasis by controlling the cholesterol synthesis through MEK/ERK signaling^37^. The study by Qijiong Li et al found that NAT10 is elevated in patients with liver cancer and the upregulated NAT10 is related to poor survival of the patients. The NAT10 protein expressions are also correlated with p53 in liver cancer tissues^38^. Yufeng Guan et al found the increased expression of HSPA1A is related to poor survival in patients with colorectal cancer^39^. By analyzing the correlation of ten single-nucleotide polymorphisms (SNPs), S P Gampenrieder et al found that EDN1 can be concerned a predictive biomarker for bevacizumab treatment in patients with metastatic breast cancer^40^. The study by Xiaohan Chang found that CACNA1C has lower expression in ovarian cancer tissues and CACNA1C could be a prognostic biomarker for patients with ovarian cancer^41^. S-Y Ma et al reported that KCNMA1 inhibits the apoptosis of ovarian cancer cells, which serves as a biomarker for patients with ovarian cancer^42^. GPCR mediates various physiologic and pathologic processes, including cancer, aging, inflammation, and diabetes dysfunctions^43–62^. The GPCR regulator β-arr1 increases liver fibrosis via Snail signaling, and β-arr1 may be a therapeutic target for liver fibrosis^63^. Moreover, the study by Yun Zhu showed that miRNA-145 inhibits the SNAI1 signaling pathway and radiation resistance in colon cancer^64^. Jinzhi Li et al found that the ELN gene expression is elevated in tumor tissues and cells. In addition, the ELN recombinant protein promotes the proliferation and progression of cancer cells^65^.

To sum up, our study identified the key signaling pathways and key biomarkers by assessing the RNA-seq data. We figured out that Alzheimer’s disease (AD) and Cushing syndrome (CS) signaling pathways were the major affected signaling pathways in hepatocellular carcinoma with the knockdown of ARHGEF2. Therefore, our study provides novel mechanisms for ARHGEF2-treated liver cancer.

## Author Contributions

Min Zhang, David C He, Andrew Chung: Methodology, Conceptualization, Writing-Reviewing and Editing.

## Funding

This work was not supported by any funding.

## Declarations of interest

There is no conflict of interest to declare.

## Notes

### Competing Interest Statement

The authors have declared no competing interest.

